# Investigating miRNA signatures in metastatic breast cancer: An *in-silico* analysis

**DOI:** 10.1101/2023.06.04.543641

**Authors:** Lakshminarasimhan Harini, Lekshmi Madhav, Sweta Srivastava, Gopalakrishna Ramaswamy, Rakesh Ramesh

## Abstract

miRNAs are small non-coding RNAs that regulate most cellular processes. Tumorigenesis disrupts the normal balance in the cell, which leads to changes in the cell cycle, cell signalling, activation of growth factors, miRNA deregulation, etc. Thus, the variations in miRNA expression between normal and tumor stages can be used to predict, diagnose, detect and identify different cancer stages, thereby suggesting the potential use of the miRNAs as biomarkers or potential therapeutic targets for cancer.

In this study we aim to identify differentially expressed miRNAs which could serve as potential biomarkers or therapeutic targets for breast cancer.

Microarray-based expression analysis was performed on tissue samples from patients with early and locally advanced breast cancer and nCounter analysis was performed to detect differentially expressed miRNA. Additionally, functional enrichment analysis, miRNA linked gene prediction, and in silico analysis for miRNA expression in cancer databases were carried out. This analysis revealed increased expression of Hsa-miR-199a and b. Further these miRs had increased expression levels across various cancer. Importantly, genes such as CALR, SSR2, and YWHAZ predicted as miR’s target were highly expressed in breast cancer. However, these genes lack therapeutic drugs targeting them. Overall, our study identified previously unexplored miRNAs and their gene targets, which could be exploited as potential targets for future therapeutic interventions.

## INTRODUCTION

MicroRNAs (miRNAs) are small, non-coding RNA molecules that play a crucial role in gene regulation [1]. These 22-nucleotide-long RNA strands are highly conserved across viruses, plants, and animals, [1] and they are involved in various cellular processes, including development, differentiation, apoptosis, and homeostasis [2]. Dysregulation of miRNAs has been linked to numerous diseases, particularly cancer. In the case of breast cancer, miRNA signatures have emerged as potential clinical markers, diagnostic tools, and therapeutic targets [3]. miRNAs are generated through a two-step cleavage process. Initially, primary microRNAs (Pri-miRNAs) are transcribed from miRNA genes and cleaved to form precursor miRNAs (pre-miRNAs). The pre-miRNAs then undergo further cleavage in the cytoplasm, resulting in the formation of mature miRNAs [4]. These mature miRNAs can regulate gene expression through post-transcriptional and translational mechanisms. They can repress translation by deadenylating or decapping the mRNA, leading to degradation. Additionally, miRNAs can upregulate transcription by binding to specific regions of the mRNA, resulting in increased expression of the target gene [5]. These regulatory mechanisms make miRNAs key players in fine-tuning gene expression and maintaining cellular homeostasis.

The dysregulation of miRNAs in breast cancer has provided a basis for utilizing miRNA signatures as therapeutic targets. By identifying specific miRNA signatures in early and late-stage breast cancer patient samples, researchers can gain insights into the underlying molecular mechanisms of metastasis. In silico analyses can be conducted to predict miRNA-regulated genes, which can then be investigated as potential therapeutic targets. Such an approach holds promise for designing targeted therapies that modulate miRNA expression or function, thereby inhibiting cancer progression and improving treatment outcomes. Understanding the miRNA signatures associated with metastatic breast cancer and identifying miRNA-regulated genes as therapeutic targets can pave the way for more personalized and effective treatments.

## METHODS

### Samples

Blood and tissue samples were collected from seven breast cancer patients across different stages including 2A, 2B, 3A, 3B and 4. The samples were collected following ethical consent obtained from the patient.

### miRNA Isolation, Quantitation and NanoString miRNA Expression assay

RNA isolation was performed using Qiagen miRNeasy Mini Kit (1038703). Tissue samples were homogenized using TOMY homogenizer in the presence of liquid nitrogen.700 µl of Qiazol was added and vortexed briefly, this mixture was incubated at RT for 5 minutes followed by the addition of 150µl of chloroform for phase separation. Samples were centrifuged at 12000*g for 15 minutes. The upper aqueous phase was transferred to a new microcentrifuge tube, and 1.5 volumes of 100% ethanol were added and mixed well by pipetting. The mixture was loaded onto the RNeasy mini column and the rest of the protocol was followed as per the manufacturer’s instructions. Elution was performed in 30µl of RNase-free water.

Quality assessment and quantitation of the RNA samples were carried out using Agilent 2100 bioanalyzer and Qubit RNA HS assay (Invitrogen Cat # Q32852). The eluted RNA was stored at -80. Quality check and quantitation of the RNA samples were performed using Agilent 2100 bioanalyzer and Qubit RNA HS assay (Invitrogen Cat # Q32852). Eluted RNA was stored at - 80. Equimolar concentrations of total RNA from different samples were pooled as per the sample details provided in **Table 1**. The samples were categorised as early breast cancer (EBC) and locally advanced breast cancer (LABC).

**Table 1:** Sample details.

| Sample# | Sample ID | Sample Name | Pool Details | Sample Type |
| --- | --- | --- | --- | --- |
| 1 | SAM1245 | DNA 4036090 | Pool 1 | **Early Breast Cancer (EBC)** |
| 2 | SAM1250 | TM 4044468<br>17/02/18 TM<br>DNA R-1 |  |  |
| 3 | SAM1260 | 4073267/<br>1335214 T<br>DNA | Pool 2 | **Locally Advanced Breast Cancer (LABC)** |
| 4 | SAM1278 | 4069989/<br>1331112<br>4069989 DNA |  |  |

For the nanoString miRNA expression assay, a 3ul aliquot was taken from the pooled sample and the assay was performed as per the nCounter miRNA Expression Assay (User Manual-MAN-C0009-07). The miRNA analysis was conducted by Theracues Innovations Pvt Ltd, Bangalore, India, using the nanoString nCounter SPRINT machine.

### miRNA expression analysis

miRNA analysis was performed on the pooled samples using the nCounter Analysis System and the nCounter Human v3a miRNA Panel which includes 798 unique clinically relevant miRNA barcodes along with appropriate positive and negative controls. The raw miRNA data in a . RCC (Reporter Code Count) format was analyzed using nSolver analysis software (NanoString technologies) version 4.0. Quality control metrics of Imaging, Binding density, positive control, and the limit of detection and ligation were checked before proceeding with the analysis. Normalization of the raw data was performed using the geometric mean of positive controls and the geometric mean of the top 100 highly expressed miRNAs as per the instructions provided in the manual.

The differential expression between the two groups (EBC and LABC) as mentioned in **Table 1**, was calculated using the build ratio module in nSolver. The EBC group was considered as the baseline for comparison. To identify the significantly differentially miRNAs, the following criteria should be fulfilled: DE call should be predicted as ‘YES’ and the geometric mean expression should be >= 40.

### In silico Analysis of the data

The fold change data (Table 2) was subjected to in silico analysis. Primarily the data were sorted based on their values and segregated into upregulated and downregulated categories. Further, FUNRICH was used to understand the molecular function, biological pathway, and biological process associated with the enriched miRNAs. Target genes for each DE miRNA were obtained from mirDB and mirTARDB. Genes exhibiting 95-100% sensitivity were chosen from miRDB while all the genes were selected from mirTARDB. These genes were further analyzed using the http://ualcan.path.uab.edu/ tool.

**Table 2.** List of top ten downregulatd miRNA and upregulated miRNA.

a) List of downregulated miRNA
| Downregulated miRNA |  |  |
| --- | --- | --- |
| miRNA |  | Fold change value |
| hsa-miR-451a | MIMAT0001631 | -70.47 |
| hsa-miR-144-3p | MIMAT0000436 | -42.15 |
| hsa-miR-522-3p | MIMAT0002868 | -9.99 |
| hsa-miR-135b-5p | MIMAT0000758 | -8 |
| hsa-miR-521 | MIMAT0002854 | -5.19 |
| hsa-miR-1299 | MIMAT0005887 | -4.68 |
| hsa-miR-514a-5p | MIMAT0022702 | -4.46 |
| hsa-miR-510-5p | MIMAT0002882 | -4.4 |
| hsa-miR-517c-3p+hsa-miR-519a-3p | MIMAT0002866 | -4.37 |

b. List of upregulated miRNA
| Upregulated miRNA |  |  |
| --- | --- | --- |
| miRNA | miRBase Accession | Fold Change |
| hsa-miR-4787-3p | MIMAT0019957 | 13.52 |
| hsa-miR-204-5p | MIMAT0000265 | 4.37 |
| hsa-miR-4454+hsa-miR-7975 | MIMAT0018976 | 3.32 |
| hsa-miR-601 | MIMAT0003269 | 2.82 |
| hsa-miR-1249-5p | MIMAT0032029 | 2.78 |
| hsa-miR-503-3p | MIMAT0022925 | 2.58 |
| hsa-miR-193a-5p+hsa-miR-193b-5p | MIMAT0004614 | 2.56 |
| hsa-miR-200b-3p | MIMAT0000318 | 2.53 |
| hsa-miR-199a-3p+hsa-miR-199b-3p | MIMAT0000232 | 2.51 |
| hsa-miR-205-5p | MIMAT0000266 | 2.51 |

Further, the STRING database was used to investigate the potential interactions between the identified genes. These same genes were also analyzed for mutation and copy number variation across breast cancer datasets from CBioPortal.

## RESULTS

### Differential expression of miRNA in LABC as compared to EBC

A comparison of the miRNA profile of EBC and LABC revealed a distinct difference between EBC and LABC. A differential gene expression analysis was performed on the two categories using nSolver. In this analysis, a negative fold change value indicates downregulation of the respective miRNA while a positive fold change value indicates upregulation. The counts in any of the samples >= 40 were considered significant. Based on these criteria, the following miRNAs were identified as differentially expressed: hsa-miR-1910-5p, hsa-miR-199a-3p, hsa-miR-199b-3p, hsa-miR-4454, hsa-miR-7975, hsa-miR-451a, hsa-miR-514a-5p, hsa-miR-522-3p, hsa-miR-1299, hsa-miR-135b-5p, hsa-miR-144-3p, hsa-miR-517c-3p, and hsa-miR-519a-3p. The miRNA was segregated based on their fold change value into downregulated and upregulated categories as listed in Table 1. Amongst these, the miRs, hsa-miR-451a, hsa-miR-144 3p, hsa-miR-522 3p, hsa-miR-135b-5p, and hsa-miR-1299 were found to be downregulated while hsa-miR-4787-3p, hsa-miR-204-5p, hsa-miR-4454+hsa-miR-7975, and hsa-miR-601 were found to exhibit positive fold change values, signifying significant upregulation.

### LABC has a distinct molecular landscape as compared to EBC

miRNAs are vital regulators of gene expression and thus participate in diverse biological processes including disease progression. In the present study, we have identified a list of miRNAs which may play an essential role in the progression of breast cancer. It is important to identify the genes regulated by these miRNAs. To analyse the genes and biological processes plausibly regulated by the altered miRNAs, we employed FUNRICH tool. FUNRICH is an open software for OMICS data analysis [6].

In terms of biological processes, the altered expression of miRNA resulted in the enrichment of several beta-oxidation pathways in LABC as compared to EBC. These include beta-oxidation of palmitoyl-CoA to myristoyl-CoA, Beta oxidation of hexanoyl-CoA to butanoyl-CoA, Beta oxidation of octanoyl-CoA to hexanoyl-CoA and Beta oxidation of decanoyl-CoA to octanoyl-CoA-CoA. These processes showed a high fold change with 386.51, 386.51, and 290.12 as respective values (Supplement data).

In the biological process category, cellular morphogenesis was highly enriched with a 196.77-fold value (Supplement data). Other enriched processes are the Transmembrane receptor protein tyrosine kinase signalling pathway, Intercellular junction assembly and/or maintenance, Negative regulation of enzyme activity, Regulation of cell shape, G-protein signalling, coupled to cyclic nucleotide second messenger, Peroxisome organization and biogenesis, Extracellular structure organization and biogenesis, Coenzyme and prosthetic group metabolism.

Several biological processes were also downregulated. These processes include Vesicle organization and biogenesis, Cellular defence response, Glycoprotein metabolism, Synaptic vesicle transport, Plasma membrane organization and biogenesis, nuclear organization and biogenesis, Reproduction, Antigen receptor-mediated signalling pathway, Xenobiotic metabolism and Neurotransmitter metabolism.

In molecular functions, Dipeptidase activity, Glucosyltransferase activity, Polysaccharide binding functions were enhanced and Sterol transporter activity, Channel or pore class transporter activity, Catalase activity, B cell receptor activity, Coreceptor activity, Oxidative phosphorylation uncoupler activity, Peptide transporter activity were downregulated.

### Expression analysis of miRNA using TCGA database

Comparison of miRNA between LABC and EBC revealed the enrichment and downregulation of some important miRNAs. We find that these miRNAs regulate important biological processes and may be thus instrumental in disease progression. We, therefore, decided to undertake a study to identify their expression in other tumours and their significance in tumour progression.

The Cancer Genome Atlas (TCGA) is a database which has genomic, transcriptomics, epigenomics and clinical data from various cancer types. It provides a vast and comprehensive dataset that enables the exploration of molecular characteristics associated with various types of cancer. OncomiR and UALCAN were thus used to analyse the TCGA dataset for expression analysis of the miRNAs chosen in this study.

OncomiR [7] provided the expression data for the chosen miRNAs across various cancer types (Fig. 2a). Amongst the DE miRNAs, hsa-miR-199a-3p and hsa-miR-199b-3p was expressed highly in almost all of the cancers except low-grade glioblastomas and ovarian cancers. These two miRNAs showed an overlap in the expression graph (Fig. 2a). The miRNA that came after them in terms of expression level was hsa-miR-451a.hsa-miR-517c-3p and hsa-miR-522-3p are expressed higher in thymus cancer and TGCT, the rare cancers on the tendon sheath. However, we did not find data for hsa-miR-1299, hsa-miR-4454+hsa-miR-7975, hsa-miR-519a-3p, hsa-miR-514a-5p, hsa-miR-4454, and hsa-miR-7975 using OncomiR. While the miRNAs, hsa-miR-1910 and hsa-miR-135b are not highly expressed in any of the cancers.

**Fig. 1:**
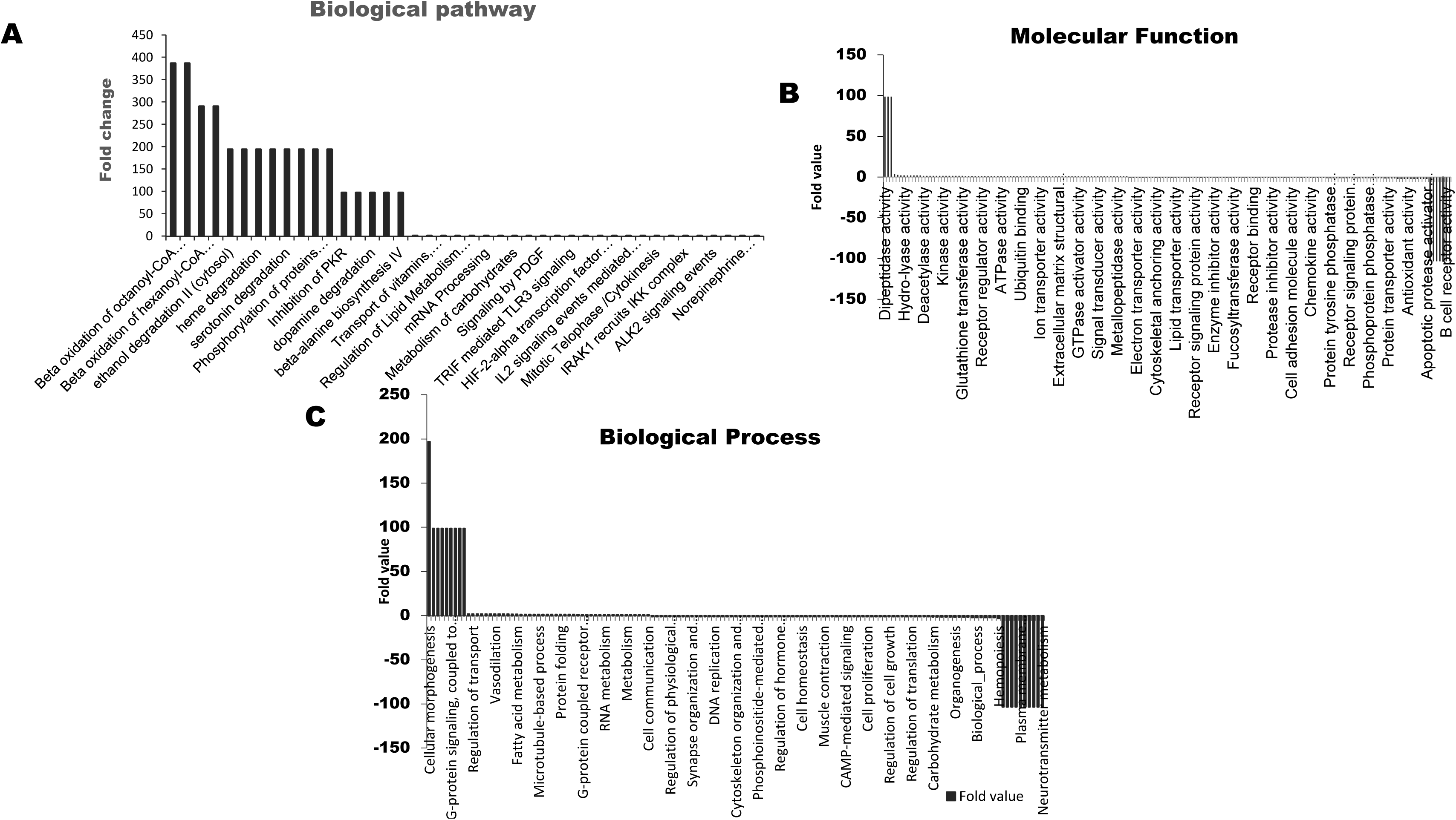
Functional enrichment of upregulated and Downregulated miRNA in FUNRICH A) Enriched beta oxidation of large and medium chain fatty acid in Biological pathway, B) Downregulated B cell receptor activity and enriched dipeptidase activity in molecular function C) Upregulated cellular morphogenesis in Biological process.

**Fig. 2:**
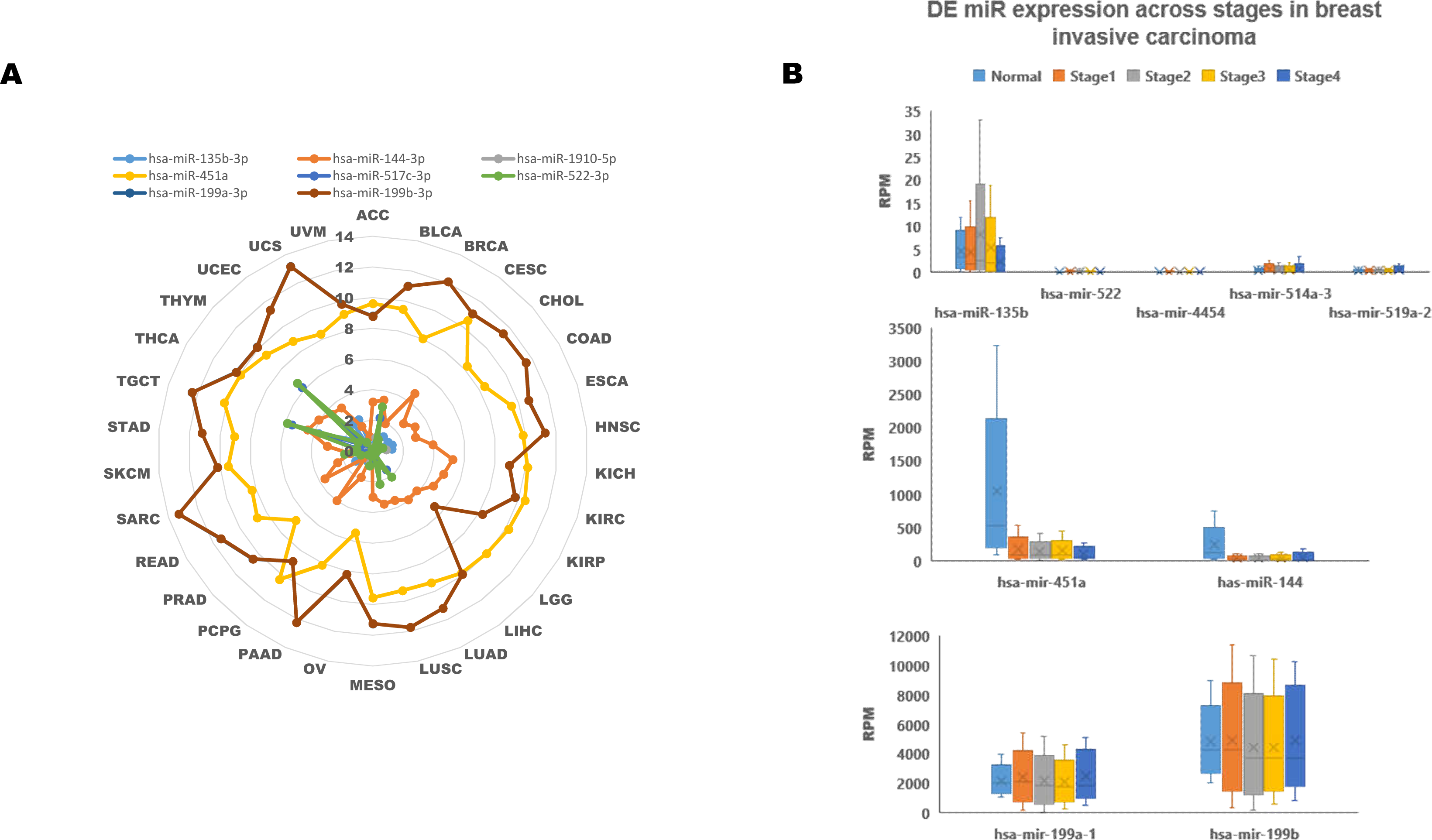
TCGA analysis of DE miRNA. A) Pan cancer analysis of DE miRNA (data from oncoMIR database) indicating the higher expression of hsa-miR-199a hsa-miR-199b and hsa-miR-451a was highly expressed in most of the cancers, B) expression of DE miRNA in different stages of breast Invasive carcinoma (data from UALCAN database). Indicates the change in expression level of miRNAs from stage I to stage IV of breast cancer patients.

A comparative analysis of these miRNAs across breast cancer stages was performed using UALCAN [8]. Figure 2b illustrates the variation in differentially expressed miRNAs among different stages of breast cancer. This analysis indicates hsa-miR-199a-3p and hsa-miR-199b-3p exhibited higher expression in cancer samples compared to the normal samples. Although a decreasing trend was observed from stages I to III, the expression of these miRNAs was slightly upregulated in stage IV. The miRs hsa-miR-144 and hsa-miR-514 were downregulated in cancer samples compared to normal. While hsa-miR135b-3p, hsa-miR-144, hsa-miR522, hsa-miR-4454, hsa-miR-451a, hsa-miR-514a-3p, and hsa-miR19a-3pwere expressed at levels lower than 500 reads per million in both cancer samples and normal samples.

### Plausible genes enriched due the miRNA

The miRNA-associated genes were identified using tools such as miRDB and mirTARBASE. miRDB contains predicted miRNA target genes based on computational algorithm while miTARBASE focuses on experimentally validated miRNA-target interactions. Thus, while using miRDB, the genes with scores greater than 95% were chosen in the analysis. However, for miTARBASE all the genes were chosen for the analysis. Table 3 and 4 presents the genes identified using miRDB and miTARBASE. Table 3 presents genes for the differentially expressed downregulated miRNAs while Table 4 presents genes for upregulated miRNA. The expression of these genes in normal and different breast cancer stages was analyzed using UALCAN and heatmaps weres generated for each gene. Fig. 3 displays the heat map of genes with significant expression level changes between normal and cancer samples MIF, YWHAZ, CALR, P4HB, TIMP1, and PSB8 were upregulated.

**Fig. 3:**
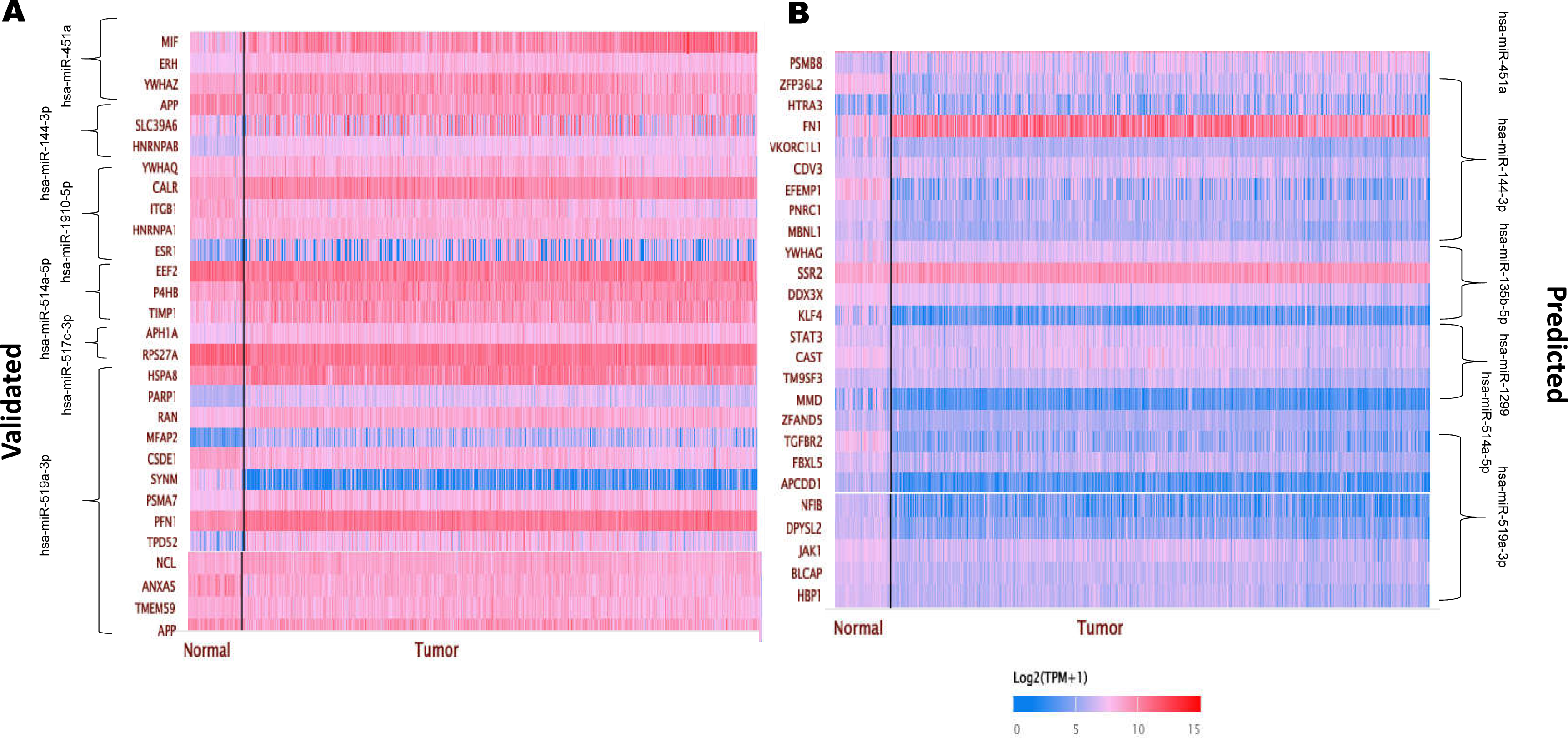
Heatmap of downregulated miRNA targeted genes expression variation in normal vs breast cancer from UALCAN database. A) Heatmap of validated upregulated target genes B) Heatmap of predicted target genes.

**Table 3:** List of selected gene targets (Predicted and Validated) for downregulated miRNA.

|  | Score | Predicted gene targets | Validated Target |  |
| --- | --- | --- | --- | --- |
| hsa-miR-135b-5p | 99 | BEND4 | hsa-miR-144-3p | PLAG1 |
|  | 99 | DOCK7 |  | FGG |
|  | 99 | LZTS1 |  | FCA |
|  | 99 | SYT2 |  | FGB |
|  | 99 | SMIM13 |  | NOTCH1 |
| hsa-miR-1299 | 100 | GPR3 | hsa-miR-1910-5p | PRRC2B |
|  | 100 | RALGAP B |  | YWHAQ |
|  | 100 | EGLN3 |  | MYL6 |
| hsa-miR-1910-3p | 98 | cAMSAP2 |  | DNAJA1 |
|  | 98 | APBB2 |  | TMEM43 |
|  | 97 | MB21D2 | hsa-miR-522-3p | SOX2 |
| hsa-miR-1910-5p | 97 | BTBD7 |  | FXN |
|  | 96 | SCN2B |  | AKAP 1 |
|  | 95 | SOS1 | hsa-miR-1299 | VAMP3 |
| hsa-miR-514a-5p | 97 | SLC22A3 |  | GNAS |
|  | 95 | STOX2 |  | SESN2 |
|  | 95 | CAMTA1 |  |  |

**Table 4:**
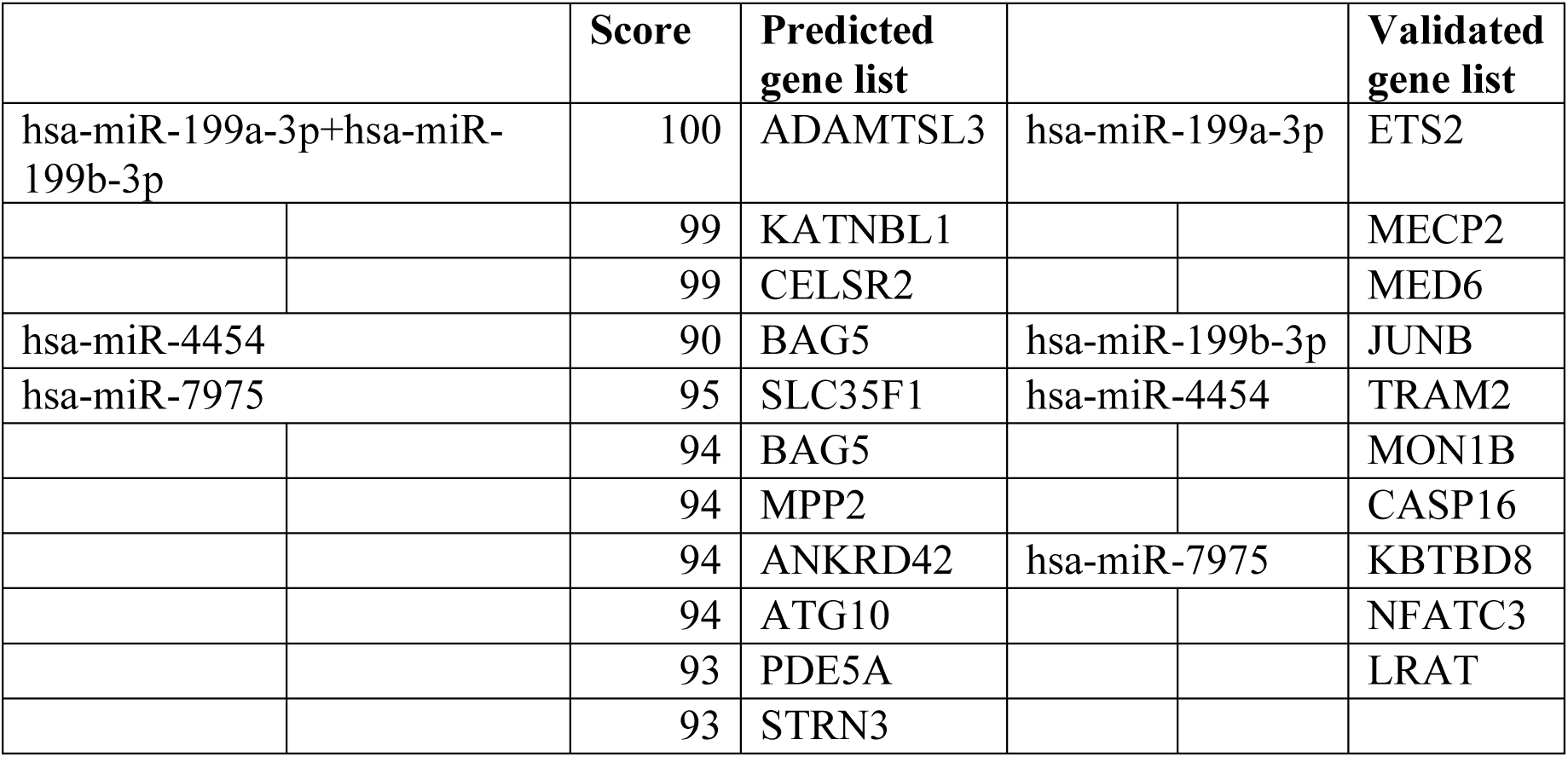
Selected list of predicted and validated gene targets for upregulated miRNA.

A further analysis of the expression of the genes between normal and cancer was performed and the genes targeted by downregulated miRs whose expression varied from normal to cancer samples are presented as listed in Table 5. Genes like MIF, YWHAZ, CALR, P4HB, TIMP1, and PSB8 were upregulated in cancer compared to normal samples. Similarly, the genes predicted to be downregulated in cancer due to overexpression of miRNAs include ETS2, JUNB, SOD2, CCDC80, and TSPAN3, and are presented in Table 5b.

**Table 5.** List of gene targets for differentially expressed miRNA.

a: List of predicted and validated target genes for the differentially expressed downregulated miRNAs.
| **miRNA** | **Validated** | **miRNA** | **Predicted** |
| --- | --- | --- | --- |
| hsa-miR-451a | MIF | hsa-miR-451a | PSMB8 |
|  | ERH | hsa-miR-144-3p | ZFP36L2 |
|  | YWHAZ |  | HTRA3 |
| hsa-miR-144-3p |  |  | FN1 |
|  | APP |  | VKORC1L1 |
|  | SLC39A6 |  | CDV3 |
|  | HNRNPAB |  | EFEMP1 |
| hsa-miR-1910-5p | YWHAQ |  | PNRC1 |
|  | CALR |  | MBNL1 |
|  | ITGB1 | hsa-miR-135b-5p | YWHAG |
|  | HNRNPA1 |  | SSR2 |
|  | ESR1 |  | DDX3X |
| hsa-miR-514a-5p | EEF2 |  | KLF4 |
|  | P4HB | hsa-miR-1299 | STAT3 |
|  | TIMP1 |  | CAST |
| hsa-miR-517c-3p |  |  | TM9SF3 |
|  | APH1A |  | MMD |
|  | RPS27A | hsa-miR-514a-5p | ZFAND5 |
| hsa-miR-519a-3p | HSPA8 | hsa-miR-519a-3p | TGFBR2 |
|  | PARP1 |  | FBXL5 |
|  | RAN |  | APCDD1 |
|  | MFAP2 |  | NFIB |
|  | CSDE1 |  | DPYSL2 |
|  | SYNM |  | JAK1 |
|  | PSMA7 |  | BLCAP |
|  | PFN1 |  | HBP1 |
|  | TPD52 |  |  |
|  | NCL |  |  |
|  | ANXA5 |  |  |
|  | TMEM59 |  |  |
|  | APP |  |  |

Table 5 (b) List of predicted and validated target genes for the differentially expressed upregulated miRNAs
| miRNA | Validated | miRNA | Predicted |
| --- | --- | --- | --- |
| hsa-miR-199a-3p+hsa-miR-199b-3p | ETS2 | hsa-miR-199a-3p | PAK4 |
|  | JUNB |  | GALNT7 |
|  | SMARCA2 | hsa-miR-7975 | TMEM183A |
|  | CAV2 |  |  |
|  | APOE |  |  |
|  | PNRC1 |  |  |
|  | QKI |  |  |
|  | IGF1 |  |  |
|  | SOD2 |  |  |
|  | HLA-B |  |  |
|  | TSPAN3 |  |  |
|  | CCDC80 |  |  |
|  | FGF1 |  |  |
|  | STAT3 |  |  |
|  | CD151 |  |  |
|  | PNRC1 |  |  |
|  | OCIAD2 |  |  |
|  | CCDC80 |  |  |
|  | SH3D19 |  |  |
|  | RBM8A |  |  |
|  | MYADM |  |  |
|  | ITPR1 |  |  |
|  | NCOA7 |  |  |

### Functional enrichment analysis of the predicted miRNA target genes to reveal the biological status of the cell

The gene lists from Table 5 a and b was further subjected to STRING analysis to predict protein-protein interactions. A full-string network with all active interaction networks and a medium confidence interaction score was chosen. Fig. 4a and 4b presents the protein interaction network for upregulated and downregulated genesets. The upregulated gene network Fig. 4a consists of 54 nodes and 153 edges with a PPI enrichment p-value of 1.0e-16. The examination of molecular functions associated with this protein network revealed that steroid hormone receptor binding, integrin binding, nuclear receptor binding, chaperone binding, nuclear hormone receptor binding, and hormone receptor binding displayed higher strength. Importantly the proteins that were involved in most of these functions were CALR, RAN, APP, APOE, FN1, ITGB1, and STAT3. The down regulated gene network consists Fig. 4b consists of 24 nodes and 13 edges with a PPI enrichment p-value of 0.0099. The examination of functions enrichment analysis for this protein network revealed Growth hormone synthesis, secretion and action has higher strength in KEGG pathways.

**Fig. 4:**
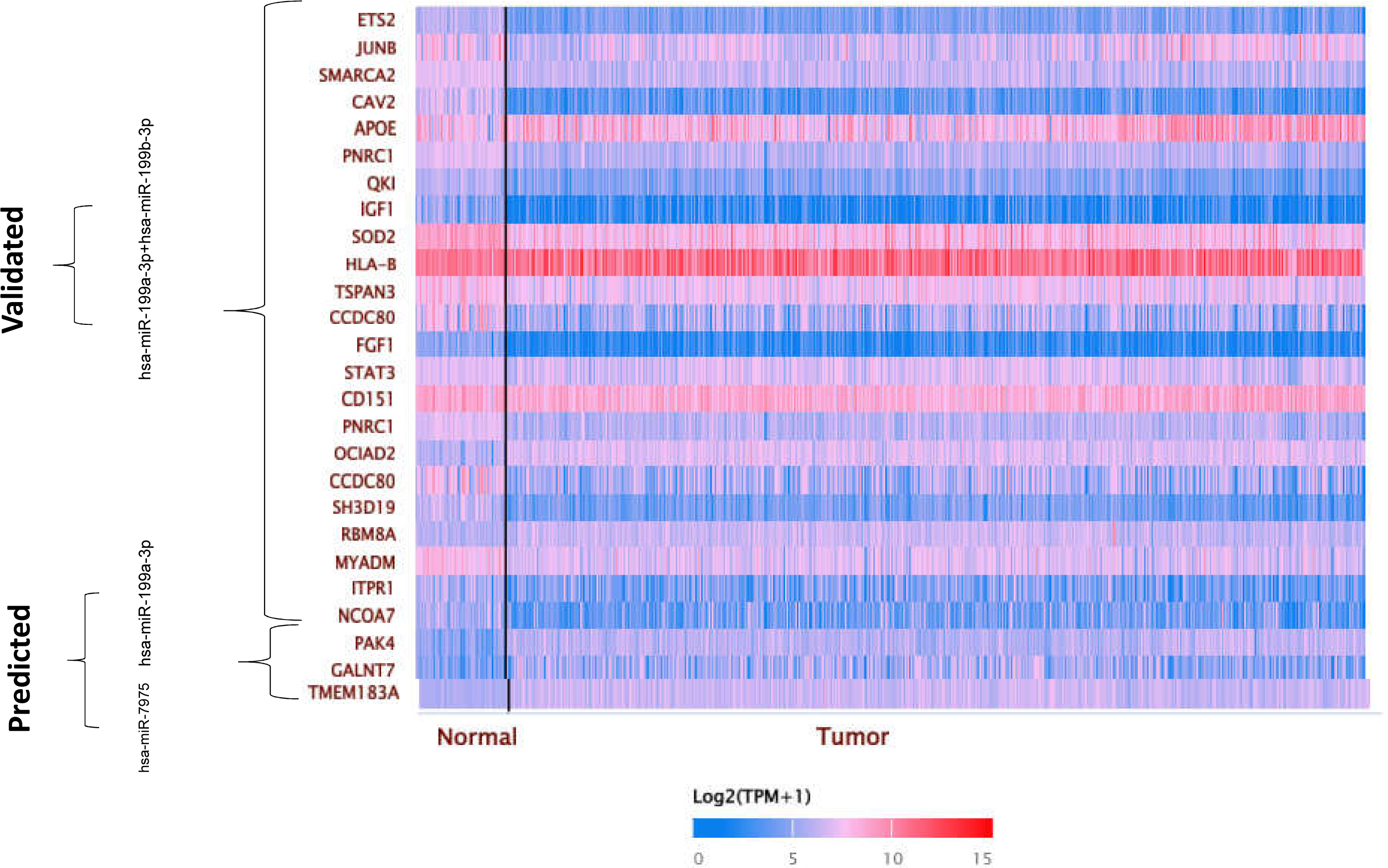
Heatmap of upregulated miRNA targeted genes expression variation in normal vs breast cancer from UALCAN database.

### Genomic variations associated with the miRNA target genes

It is important to note that genetic variations within the miRNA target genes can affect miRNA binding sites and alter gene expression levels. Single Nucleotide Polymorphisms, insertions, deletions, copy number variations and other structural variations can affect gene expression patterns. Thus, an analysis of the genomic variation was undertaken for the genes listed in Tables 5a and 5b.

Mutation and copy number variation in the genes was analysed using CBioPortal. The analysis focussed on different types of breast cancer, including invasive breast carcinoma, adenoid cystic breast carcinoma (2015), and metastatic breast cancer 2021. The data was from studies from MSK cancer cells 2018, MSK nature cancer 2022, MSK cancer discovery 2022, HTAN2022, MSK clinical cancer research 2022, MSK NPJ breast cancer 2019, SMC 2018, proteogenomic landscape breast cancer CPTAC 2020, and metastatic breast cancer project 2020. A total of 7063 samples were queried. The analysis revealed that gene amplification is the major alteration observed in most of the genes. TMEM183A exhibited the highest amplification rate of 25%. SSR2 and YWHAZ genes also displayed significant amplification rates, with 21% and 19%, respectively. Other genes, such as APH1A (18%), RBM8A (17%), TPD52 (17%), and PARP1 (14%), were also found to be amplified in the samples.

### Drug target prediction miRNA-associated genes

It is imperative to study the possibility of therapeutic possibility using the identified targets. To explore this possibility, a study was performed to identify drugs targeting the genes associated with miRNA expression. Drug availability for CALR, RAN, APP, APOE, FN1, ITGB1, STAT3, SSR2, TMEM183A, RBM8A, APH1A, TPD52, and PARP was retrieved from the Therapeutic Target Database [9]. Table 6 lists the drugs for these targets. It is to be noted that not all the genes identified in this study have a drug suited for therapy, however, APOE, FN1, ITGB1 and STAT3 show a targeted drug available.

**Table 6:** List of genes and its available drugs from Drug Target Database.

| Target | Drug |
| --- | --- |
| CALR | No drug available |
| RAN | No information |
| APP | No drug available |
| APOE | AEM-28 |
| FN1 | Ocriplasmin |
| ITGB1 | 131I-radretumab |
| STAT3 | Acitretin, ISIS-STAT3 |
| SSR2 | No information |
| TMEM183A | No information |
| RBM8A | No information |
| APH1A | No information |
| TPD52 | No information |
| PARP | Nicotinamide |
| YWHAZ | No information |

The availability of targeted drugs for these genes opens possibilities for further research and development of therapies that aim to modulate their expression or function. Overall, the study provides valuable insights into potential therapeutic targets and highlights the importance of ongoing research and development to uncover new drugs and therapies for various conditions associated with miRNA expression.

## DISCUSSION

miRNA (microRNA) analysis is an important tool in cancer research and diagnosis [10]. They are small non-coding RNA molecules and play a significant role in gene regulation by binding to messenger RNA (mRNA) and consequently regulating various cellular processes such as cell growth and proliferation. miRNA expression variation can be used as Biomarkers for cancer diagnosis, subtyping of tumours, prognostic indicator, and therapeutic targets [11]. In the present study, miRNA analysis was conducted to understand the difference between LABC and EBC with respect to miRNA expression.

A comparison of upregulated and downregulated miRNAs reveals enhancement of long and medium-chain fatty acid beta-oxidation, changes in cellular morphogenesis, and downregulation of B cell-mediated immune reaction. These findings correlate with previous studies indicating that beta-oxidation of fatty acids in cancer relates to proliferation, survival, stemness, drug resistance, and metastatic progression, which drives breast cancer cells to EMT leading to metastasis [12, 13]. Another important aspect of cancer progression is alteration in immune response. Enrichment analysis of EBC vs. LABC in biological processes and pathways confirms the downregulation of B cell receptor activity and cellular defence mechanisms. Recent studies on the immune system adaptation of metastatic cancer cells have discovered that these cells adapt to or evade the immune response and survive [14].

A TCGA based analysis of the expression patterns of the identified miRNAs revealed that hsa-miR-199a and -b are highly upregulated in most cancers compared to normal samples while hsa-miR-451a and hsa-miR-144-3p are downregulated in breast cancers compared to normal cells.

Whereas, Results from the miRBase and RNA databases suggest that hsa-miR-199a along with hsa-miR-199b negatively regulate matrix metalloproteinase and positively regulate endothelial migration. Another study on hsa-miR-199a-3p suggests its targeting of BRCA gene and sensitisation of Triple negative breast cancer (TNBC) to therapy [15]. We also observe that hsa-miR-451a was significantly downregulated in the LABC sample. Hsa-miR-451a has been reported to target CCND2, contributing to breast cancer progression [16]. Another miRNA that was downregulated in LABC tissue was hsa-miR-144a. Downregulation of hsa-miR-144a correlates to a poor prognosis in colorectal cancers [17].

Analysis of genes targeted by the differentially expressed miRNAs revealed CALR, RAN, APP, APOE, FN1, ITGB1, and STAT3 as plausible targets. These genes play a key role in cancer progression. Interestingly, APOE, FN1, ITGB1 and STAT3 have available inhibitors and can be used for treatment of cancers. Calreticulin protein is involved in the invasiveness and metastasis of breast cancer [18] and has been identified as a potential biomarker in various cancers [19]. However, there isn’t a good inhibitor of CALR which can be used as a therapeutic molecule. Similarly, RAN is also a biomarker for cancer progression and overexpression of RAN indicates invasiveness and metastasis in breast cancer [20]. Unfortunately, similar to CALR, there is no effective inhibitor for RAN. Our analysis also revealed that genes including SSR2, TMEM183A, RBM8A, YWHAZ, APH1A, PARP1, and TPD52 showed higher genetic alterations. SSR2 is an endoplasmic reticulum transmembrane protein that plays a role in ER stress in HCC patients [21], however, its role in breast cancers is yet to be explored. Additionally, there is no available information on SSR2 in the drug target database.

Analysis reveals that the most downregulated miRNA in advanced cancers is hsa-miR-451a and it regulates the expression of genes including PSMB8, MIF, ERH, and YWHAZ. STRING analyses of the gene set revealed that YWHAZ, a transmembrane protein, is a common node for mitochondrial matrix permeabilization and transcriptional regulation. as it interacts with proteins like PAK4, YWHAG, YWHAQ, PFN1, ITGB, etc. Interestingly, the CBioPortal analysis Fig. 6 indicates the YWHAZ gene has a higher level of genetic alteration.

**Fig. 5:**
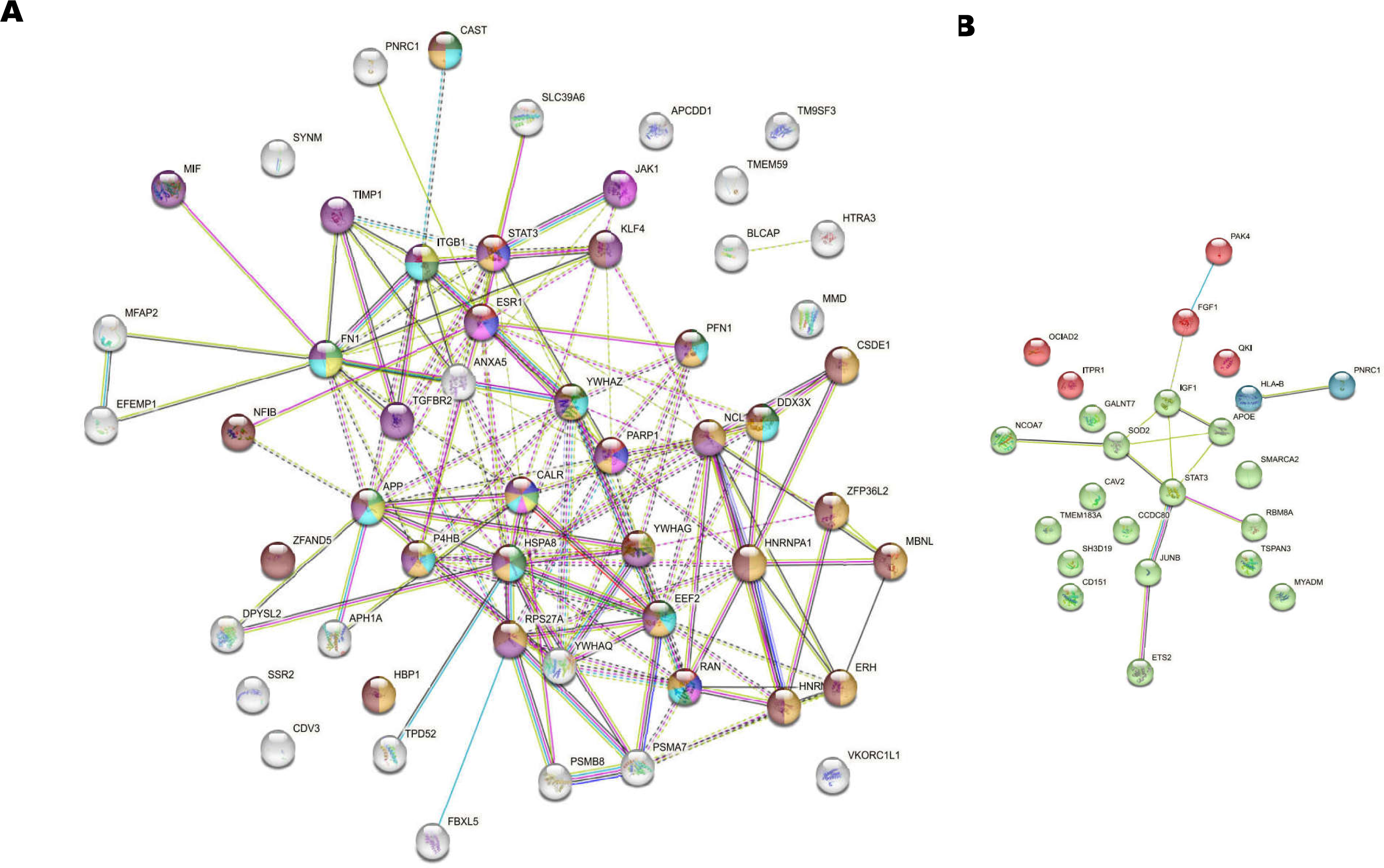
STRING interaction of genes A) upregulated gene targets and B) downregulated gene targets selected based on expression analysis from UALCAN. Genes like YWHAZ, RAN, ITGB1, STAT3, FN1 etc were found to be the common genes involved in most of the molecular function in enrichment analysis.

**Fig. 6:**
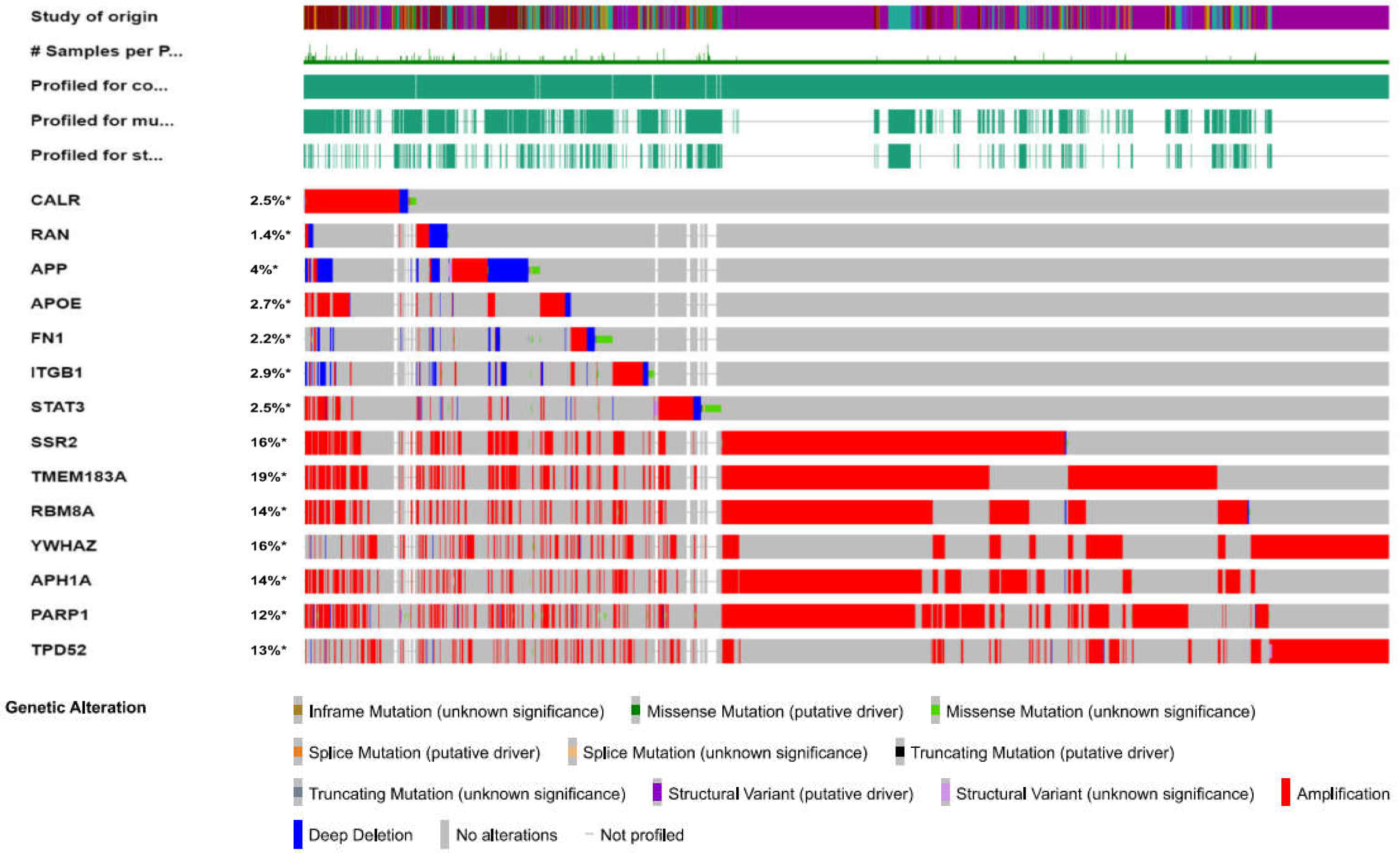
Genetic alteration in target genes: CBioPortal images of mutations and amplification of selected genes from STRING interactions in breast cancer patients.

The miRNA signature identified from the metastatic breast cancer patient tissues in this study shows that hsa-miR199a/b are upregulated while hsa-miR-451a/3p and hsa-miR-144-3p are downregulated.

The availability of targeted drugs for some genes opens possibilities for further research and development of therapies that aim to modulate their expression or function. Additionally, the study identified novel targets to develop novel inhibitors. Overall, the study provides valuable insights into potential therapeutic targets and highlights the importance of ongoing research and development to uncover new drugs and therapies for various conditions associated with miRNA expression.

## CONCLUSION

This study identifies miRNA signatures in early and late breast cancer tissue. This was further scrutinized *in silico* for the prediction of the potential therapeutic target. The role of proteins CALR, SSR2 and YWHAZ could be further explored as a therapeutic target.

## Reference

1. Macfarlane, Leigh-Ann, and Paul R Murphy. MicroRNA: Biogenesis, Function and Role in Cancer. Current genomics. 11,7 (2010): 537-61. doi:10.2174/138920210793175895

2. Gebert, L.F.R., MacRae, I.J. Regulation of microRNA function in animals. Nat Rev Mol Cell Biol 20, 21–37 (2019). 10.1038/s41580-018-0045-7.

3. Long Xinghua, Shi Yu, Ye Peng, Guo Juan, Zhou Qian, Tang Yueting MicroRNA-99a Suppresses Breast Cancer Progression by Targeting FGFR3, Frontiers in Oncology, 9, (2020). Doi:10.3389/fonc.2019.01473.

4. O’Brien Jacob, Hayder Heyam, Zayed Yara, Peng Chun. Overview of MicroRNA Biogenesis, Mechanisms of Actions, and Circulation, Frontiers in Endocrinology, 9, (2018). doi: 10.3389/fendo.2018.00402.

5. JOUR, Chen, Shen Liang, Valinezhad Orang, Ayla; Safaralizadeh, Reza; Kazemzadeh-Bavili, Mina. 2014, Mechanisms of miRNA-Mediated Gene Regulation from Common Downregulation to mRNA-Specific Upregulation, International Journal of Genomics (2014) doi: 10.1155/2014/970607.

6. Pamali Fonseka Mohashin Pathan Sai V.Chitti Tae youngKang Suresh Mathivanan, FunRich enables enrichment analysis of OMICs datasets, Journal of Molecular Biology 433, 11, (2021) 166747.

7. Wong N, Chen Y, Chen S, Wang X OncomiR: an online resource for exploring pan-cancer microRNA dysregulation. Bioinformatics, 34(4) (2018) 713–715.

8. Chandrashekar DS, Bashel B, Balasubramanya SAH, Creighton CJ, Rodriguez IP, Chakravarthi BVSK and Varambally S. UALCAN: A portal for facilitating tumor subgroup gene expression and survival analyses. Neoplasia. (2017) :649–658. doi: 10.1016/j.neo.2017.05.002.

9. Y. Zhou, Y. T. Zhang, X. C. Lian, F. C. Li, C. X. Wang, F. Zhu, Y. Q. Qiu and Y. Z. Chen. Therapeutic target database update 2022: facilitating drug discovery with enriched comparative data of targeted agents. Nucleic Acids Research. 50(D1): (2022) 1398–1407.

10. Galvão-Lima, L.J., Morais, A.H.F., Valentim, R.A.M., et al. miRNAs as biomarkers for early cancer detection and their application in the development of new diagnostic tools. BioMed Eng OnLine 20, 21 (2021). doi: 10.1186/s12938-021-00857-9

11. Bertoli G, Cava C, Castiglioni I. MicroRNAs: New Biomarkers for Diagnosis, Prognosis, Therapy Prediction and Therapeutic Tools for Breast Cancer. Theranostics. 13 5(10) (2015) 1122–43. doi: 10.7150/thno.11543.

12. Ma Y, Temkin SM, Hawkridge AM, Guo C, Wang W, Wang XY, Fang X. Fatty acid oxidation: An emerging facet of metabolic transformation in cancer. Cancer Lett. 28;435 (2018) 92–100. doi: 10.1016/j.canlet.2018.08.006.

13. Loo SY, Toh LP, Xie WH, Pathak E, Tan W, Ma S, Lee MY, Shatishwaran S, Yeo JZZ, Yuan J, Ho YY, Peh EKL, Muniandy M, Torta F, Chan J, Tan TJ, Sim Y, Tan V, Tan B, Madhukumar P, Yong WS, Ong KW, Wong CY, Tan PH, Yap YS, Deng LW, Dent R, Foo R, Wenk MR, Lee SC, Ho YS, Lim EH, Tam WL. Fatty acid oxidation is a druggable gateway regulating cellular plasticity for driving metastasis in breast cancer. Sci Adv. 7(41) (2021). doi: 10.1126/sciadv.abh2443.

14. Blomberg OS, Spagnuolo L, de Visser KE. Immune regulation of metastasis: mechanistic insights and therapeutic opportunities. Dis Model Mech. 11(10) (2018). doi: 10.1242/dmm.036236

15. Ho JC, Chen J, Cheuk IW, Siu MT, Shin VY, Kwong A. MicroRNA-199a-3p promotes drug sensitivity in triple negative breast cancer by down-regulation of BRCA1. Am J Transl Res. 14(3) (2022),:2021–2036.

16. Zhang H, Chen P, Yang J. miR-451a suppresses the development of breast cancer via targeted inhibition of CCND2. Mol Cell Probes. 54:101651. (2020) doi: 10.1016/j.mcp.2020.101651.

17. Li T, Tang C, Huang Z, Yang L, Dai H, Tang B, Xiao B, Li J, Lei X. miR-144-3p inhibited the growth, metastasis and epithelial-mesenchymal transition of colorectal adenocarcinoma by targeting ZEB1/2. Aging (Albany NY). 13(13) (2021) 17349–17369. doi: 10.18632/aging.203225.

18. Zamanian, M., Qader Hamadneh, L.A., Veerakumarasivam, A. et al. Calreticulin mediates an invasive breast cancer phenotype through the transcriptional dysregulation of p53 and MAPK pathways. Cancer Cell Int 16, 56 (2016). 10.1186/s12935-016-0329-y

19. Li Y, Liu X, Chen H, Xie P, Ma R, He J, et al. Bioinformatics analysis for the role of CALR in human cancers. PLoS ONE 16 (12), (2021) e0261254. Doi: 10.1371/journal.pone.0261254.

20. Boudhraa Z, Carmona E, Provencher D, Mes-Masson AM. Ran GTPase: A Key Player in Tumor Progression and Metastasis. Front Cell Dev Biol. 8, 345 (2020). doi: 10.3389/fcell.2020.00345.

21. Zhang G, Sun J. Endoplasmic Reticulum Stress-Related Signature for Predicting Prognosis and Immune Features in Hepatocellular Carcinoma. J Immunol Res. (2022) 1366508. doi: 10.1155/2022/1366508.

